# EEG Functional Connectivity as a Marker of Evolution from Infantile Epileptic Spasms Syndrome to Lennox-Gastaut Syndrome

**DOI:** 10.1101/2025.04.30.650531

**Authors:** Blanca Romero Milà, Virginia B. Liu, Rachel J. Smith, Derek K. Hu, Natalie A. Benneian, Shaun A. Hussain, Maija Steenari, Donald Phillips, David Adams, Clare Skora, Beth A. Lopour, Daniel W. Shrey

**Affiliations:** Department of Biomedical Engineering, University of California, Irvine, Irvine, CA, United States; Division of Neurology, Children’s Hospital Orange County, Orange, CA, United States; Department of Pediatrics, University of California, Irvine, Irvine, CA, United States; Department of Electrical and Computer Engineering, University of Alabama at Birmingham, Birmingham, AL, United States; Department of Biomedical Engineering, California State University, Long Beach, Long Beach, CA, United States; Department of Computer Engineering and Computer Science, California State University, Long Beach, Long Beach, CA, United States; Division of Pediatric Neurology, University of California, Los Angeles, CA, United States

**Author notes:** Daniel W. Shrey and Beth A. Lopour contributed equally to this work.

## Abstract

**Background and Objectives:** Timely diagnosis and effective treatment of Lennox-Gastaut Syndrome (LGS) improve prognosis and lower healthcare costs, but the transition from infantile epileptic spasms syndrome (IESS) to LGS is highly variable and insidious. Objective biomarkers are needed to monitor this progression and guide clinical decision making.

**Methods:** We retrospectively collected longitudinal EEG data at the Children’s Hospital of Orange County from fifteen children who were diagnosed with IESS and later with LGS between 2012 and 2021. EEGs were from IESS and LGS diagnoses, between the two diagnoses, and following LGS diagnosis. Functional connectivity networks were calculated using a cross-correlation-based method and assessed relative to diagnostic timepoint, treatment response, presence of clinical markers of disease, age, and amplitude of interictal spikes.

**Results:** Connectivity strength was high at LGS diagnosis and decreased after favorable response to treatment, but it remained stable or increased when response was unfavorable. In all subjects, connectivity strength was higher at the time of LGS diagnosis than at the preceding timepoint. Presence of clinical markers of LGS were associated with high connectivity strength, but no single marker predicted connectivity strength.

**Discussion:** Computational EEG analysis can be used to map the evolution from IESS to LGS. Changes in connectivity may enable prediction of impending LGS and treatment response monitoring, thus facilitating earlier LGS treatment and guiding medical management.

**Key points:**

- EEG functional connectivity analysis can track progression from infantile epileptic spasms syndrome (IESS) to Lennox-Gastaut Syndrome (LGS).
- High connectivity strength at LGS diagnosis decreases with favorable treatment response but remains high with poor response.
- Clinical LGS markers correlate with high connectivity, but no single marker predicts connectivity strength.
- EEG functional connectivity analysis may help predict LGS onset, enabling early intervention and improving prognosis.

## 1. Introduction

Lennox-Gastaut Syndrome (LGS) is a severe epilepsy characterized by a triad of encephalopathy, multiple seizure types, and electroencephalogram (EEG) findings of slow spike-and-wave (SSW) and generalized paroxysmal fast activity (GPFA) [1]. For patients with LGS, long-term neurocognitive outcomes are frequently poor, and complete seizure freedom is rarely achieved, often impacting social function, the likelihood of independence, and quality of life. LGS patients are often treated with multiple anti-seizure medications, yet the decrease in seizure frequency due to adding a new medication to their regimen is typically only 15% to 26% [2-4]. Timely diagnosis and effective treatment of LGS can significantly improve outcomes and reduce disease management costs [5]. However, the diagnosis of LGS relies on a combination of EEG features that can vary across patients and time, are potentially altered by ongoing treatments, and are common to other epilepsy syndromes [1].

Further confounding LGS diagnosis is the fact that 20-40% of children with LGS have a history of infantile epileptic spasms syndrome (IESS) [1, 6, 7]. IESS is an epileptic encephalopathy that occurs in children younger than two-years-old. This syndrome is associated with epileptic spasms occurring in clusters, often with a characteristic interictal EEG finding called hypsarrhythmia. IESS and LGS have overlapping features, and the transition between them is gradual, lasting several months to several years or more [7]. Patients who evolve from IESS to LGS are less likely to respond favorably to medication compared to those with LGS and no history of IESS [8]. The only currently known predictive clinical factor that portends evolution from IESS to LGS is having an underlying brain disease [7]. No relationship has been found between LGS and specific etiologies or the age of spasms onset [6]. Therefore, there is a need for robust biomarkers to monitor the progression from IESS to LGS and guide treatment decisions.

Previous studies have primarily focused on clinical data, such as treatment outcome and risk factors [9]; however, computational EEG analysis can provide a means to objectively assess diagnostic criteria, predict initial medication response, and monitor disease evolution. For example, EEG functional connectivity networks (FCNs) have shown promise as a biomarker of IESS. Specifically, cross-correlation FCNs have been shown to be stronger in children with IESS compared to controls during wakefulness and sleep [10-12]. These networks have been found to be individualized rather than stereotyped within a subject group [13], and they demonstrate high test-retest reliability, as the cumulative calculated FCN stabilizes with as little as 150 seconds of EEG data [11, 12, 14]. Additionally, changes in cross-correlation FCNs reflect treatment response in children with IESS treated with ACTH and/or vigabatrin as first-line therapy. In a study of 21 children with IESS, decreased FCN strength following IESS treatment was correlated with favorable clinical treatment response [12].

Therefore, we hypothesize that EEG-based functional connectivity can serve as a robust, objective biomarker for mapping the evolution from IESS to LGS and for evaluating treatment response. To our knowledge, this is the first study to explore computational EEG biomarkers during the transition from IESS to LGS. Given how little is known about this dynamic period, our research is a crucial first step toward developing objective biomarkers which could significantly improve our understanding and management of epilepsy in this population.

## 2. Methods

### 2.1. Data Characteristics

Approval to perform this retrospective observational study was granted by the Children’s Hospital of Orange County (CHOC) Institutional Review Board. We collected longitudinal scalp EEG data during sleep from fifteen subjects diagnosed with IESS and, subsequently, LGS at CHOC between 2012 and 2021. As inclusion criteria, subjects had both IESS and LGS diagnostic EEGs done at CHOC with at least one EEG recorded during the interim between the diagnoses (Table 1). Subjects were identified by querying the electronic medical record using ICD-9/10 codes for infantile spasms (345.60-61, G40.821-824) and LGS (345.11, 345.80-81, G40.811-814) then confirming diagnoses via chart review. The LGS diagnostic timepoint was defined cumulatively; we required that the subject exhibited SSW activity along with at least one characteristic LGS seizure type at some point during their care. We did not require that SSW and multiple seizure types be captured all in a single EEG recording.

**Table 1:**
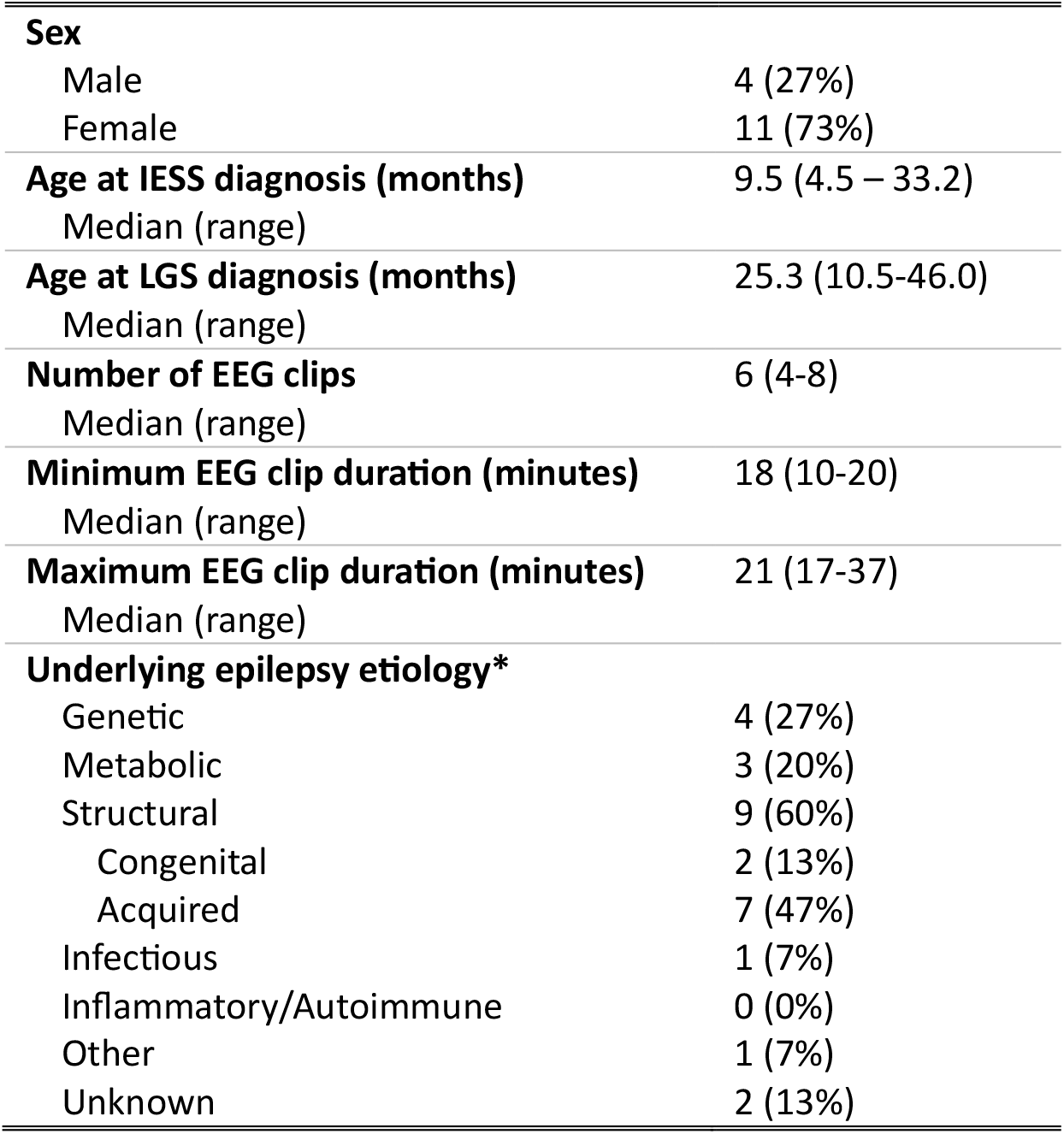
Subject and EEG data characteristics. *Note that some subjects had underlying etiologies with multiple classifications, e.g. trisomy 21 and hypoxic-ischemic encephalopathy, hence the total number of etiologies (n=20) exceeds the number of subjects in the study (n=15).

We analyzed EEG clips from multiple timepoints, including IESS diagnosis, LGS diagnosis, and a maximum of five interim timepoints between the diagnoses. If more than five interim EEG recordings were available, we selected the subset of five in which the inter-EEG time intervals were as equal as possible. We also included the first EEG acquired after LGS diagnosis and treatment, which was available for all subjects except subject 12. The EEG data, which included both routine studies and overnight studies, were recorded from nineteen scalp electrodes using the International 10-20 system, including FP1, FP2, F3, F4, C3, C4, P3, P4, O1, O2, F7, F8, T3, T4, T5, T6, Fz, Cz, and Pz, sampled at 200 Hz.

### 2.2. Data pre-processing

EEG clips consisted of the first 20 minutes of definitive non-REM sleep in the recording. Each clip was re-referenced to the common average, artifacts were identified using an amplitude-based automatic artifact detector [10], and data were filtered with a zero-phase broadband filter (0.5 – 55 Hz). Then clips were windowed into one-second epochs with no overlap, and 600 epochs without artifacts were randomly selected from each EEG from every patient and used to calculate functional connectivity.

### 2.3. Connectivity calculation: Cross-correlation

Connectivity was calculated using the cross-correlation method previously applied to data from both healthy subjects [16, 17] and those with IESS [10]. Prior studies have established this as a robust and reliable method for analysis of infant EEG, with high potential for clinical relevance as a computational biomarker of IESS [10, 12, 18]. The method was first described in Kramer et al., and Chu et al. [16, 19]; we briefly summarize it here. First, each one-second epoch of EEG was normalized to have zero mean and unit variance. Then, we calculated the absolute value of the cross-correlation (maximum lag of 200 ms) for each electrode pair and epoch and identified the maximum cross-correlation magnitude [19]. This value was subsequently normalized by its autocorrelation. Next, significance was assessed by comparing the resulting normalized value to a baseline distribution generated under the null hypothesis that there was no temporal relationship between each pair of signals. This baseline distribution was estimated by shufling the 1-second epochs of data in time and then calculating the normalized maximum cross-correlation between all channels, as previously described. This process was repeated 1000 times, and the resulting correlation values were sorted to generate a cumulative baseline distribution. For each epoch, the connection between two EEG channels was considered significant when the maximum cross-correlation exceeded the 95^th^ percentile of the baseline distribution [12].

To eliminate spurious connectivity resulting from volume conduction, epochs where the max cross-correlation occurred at zero time lag were deemed nonsignificant [16]. Forward modeling simulations demonstrated that this method successfully identified all spurious connections attributed to volume conduction; while this is thought to be a conservative approach, it was reported that its implementation does not dramatically alter the final connectivity network, suggesting that it is not overly conservative [16]. Connectivity strength of each electrode pair was defined as the fraction of one-second epochs with significant cross-correlation. Thus, a connection with a strength of 0.25 had significant cross-correlation values in 25% of one-second epochs. Finally, we quantified the overall connectivity strength of the network in two ways: (1) the number of connections with connectivity strength greater than 0.1 and (2) the mean connectivity strength calculated using the 10% of connections with the highest connectivity strength values. The threshold of 0.1 was used to define “strong” connections, as a previous study demonstrated that this approach produces robust results across thresholds ranging from 0.09 to 0.23 [12]. Overall, we found that these two methods were equivalent, as the values of these two metrics were highly correlated (Pearson test, p-value < 0.001, R = 0.94). Therefore, only the number of strong connections is reported here.

### 2.4. Calculation of interictal epileptiform discharge (IED) amplitude

Interictal epileptiform discharges (IEDs) may occur frequently in the EEGs of patients with IESS or LGS, and they can have high amplitudes that dominate the signal. Because this could significantly impact the connectivity measurement, we tested the correlation between IED amplitude and connectivity strength. IEDs were detected in each EEG clip using Persyst® software, and the mean spike amplitude for each clip was calculated. Persyst® specified the channel closest to the source of the majority of IEDs in each patient, and a custom MATLAB script analyzed the EEG from this channel to calculate the amplitude as follows.

Following the American Clinical Neurophysiology Society (ACNS) guide, data were referenced using a longitudinal bipolar montage, and IED amplitude was measured from peak to trough using the highest and lowest local extrema in a 500 ms window surrounding each detected IED (+/-250 ms). The EEG in each window was required to have both a positive and a negative peak; otherwise, it was disregarded, as the detection was assumed to be a false positive. The window length of 500 ms was chosen to capture the entire IED waveform, as IED duration can vary from 9 to 200 ms, with a subsequent deflection lasting between 130 and 200 ms. Finally, the relationship between spike amplitude and connectivity strength for each EEG recording was assessed using the Pearson correlation coefficient.

### 2.5. Visual EEG analysis

Each EEG was visually analyzed by board-certified pediatric epileptologists to determine disease state based on standardized clinical markers. At IESS diagnosis, patients had epileptic spasms with abnormal EEG background, with or without hypsarrhythmia. Subjects 4 and 10 were marked as not having spasms at the first EEG because, although they presented clinically with spasms, these events were not captured during the initial EEG study. After treatment, “ES-” was used to indicate those with no clinical spasms at the time of the EEG; those with persistent spasms were designated “ES+”. LGS diagnosis was defined as the first timepoint at which the standard diagnostic triad of LGS had cumulatively been observed, with LGS EEG features occurring simultaneously in one EEG or over multiple EEG studies. Following LGS treatment, “SSW+” (or “SSW-”) was used to indicate the presence (absence) of the slow spike-and-wave EEG pattern. The presence or absence of ongoing encephalopathy and uncontrolled seizures could not be robustly assessed due to the study’s retrospective nature. Further, each EEG was assessed for the presence or absence of hypsarrhythmia, epileptic spasms, GPFA, and SSW during the entire EEG study. The epileptologist was blinded to the patient’s clinical state. Note that only clips (subsets) of the full EEG studies were computationally analyzed, and there was no requirement that these features be present in the clips. In fact, none of the EEG clips contained epileptic spasms. This approach ensures that EEG connectivity is an effective marker when applied to a patient’s baseline background EEG, rather than requiring direct capture of ictal EEG.

## 3. Results

### 3.1. Connectivity strength is high at LGS diagnosis and decreases with SSW resolution

In all subjects, connectivity strength was higher at the time of LGS diagnosis compared to the preceding EEG (Figure 1A, Wilcoxon signed-rank sum test, p < 0.05).

**Figure 1:**
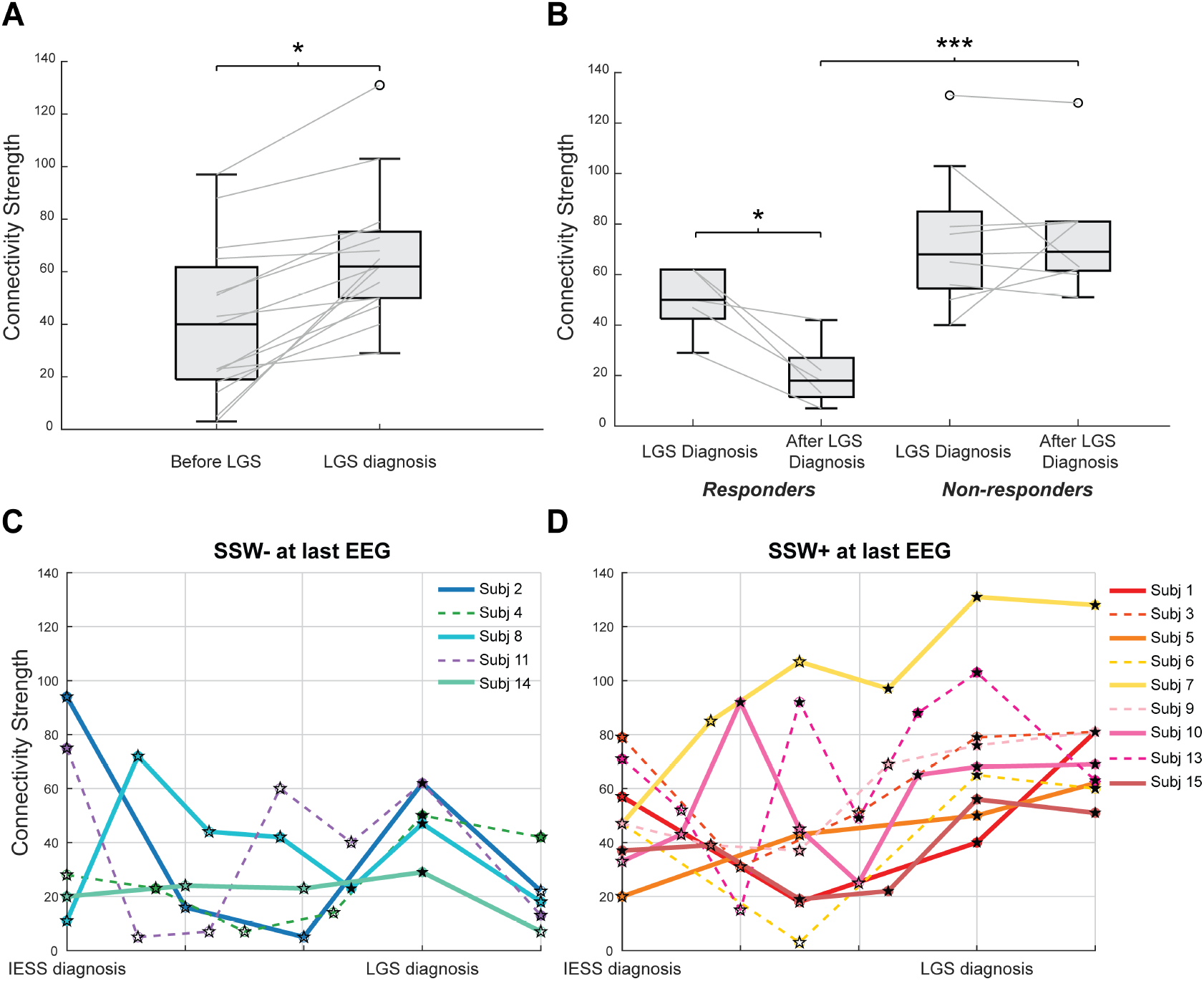
Connectivity strength is a candidate biomarker for LGS diagnosis and treatment response. (A) EEG connectivity strength is higher at the time of LGS diagnosis (right) than at the preceding time point (left, Wilcoxon signed-rank sum test, p < 0.05). (B) EEG connectivity strength decreases after LGS treatment for subjects with resolution of SSW after treatment (left), while changes were not consistent for subjects with non-resolution of SSW (right). Data are shown for the EEG at LGS diagnosis and the subsequent timepoint. Wilcoxon rank sum test, *: p < 0.05; ***: p < 0.001. (C) Subjects who ultimately demonstrated resolution of SSW generally had connectivity strength < 60 before and after LGS diagnosis. (D) Subjects who ultimately demonstrated persistent SSW exhibited a trend of increasing connectivity strength over time, generally ending with connectivity strength > 60 during their final EEG. For subfigures (C) and (D), the x-axis shows normalized time, where the times of IESS and LGS diagnoses are aligned for all subjects. Note that subject 12 was excluded from this analysis due to a lack of EEG recordings after LGS treatment. Each colored line represents longitudinal data from one subject. The markers indicate the presence of clinical markers, with circles for ES and stars for SSW. Further, filled and open markers correspond to ES+/SSW+ and ES-/SSW-, respectively.

We then compared the LGS diagnosis EEG to the subsequent EEG, recorded following LGS-specific treatment initiation. All subjects with post-treatment resolution of the SSW pattern (responders) demonstrated a decrease in connectivity strength (Wilcoxon rank sum test, p < 0.05), whereas there was no significant change in connectivity for those who had persistent SSW (Figure 1B). We observed the same trend with IESS; after treatment, subjects with resolution of IESS clinical markers had significant decreases in connectivity strength (Wilcoxon rank sum test, p < 0.01), but non-responders exhibited no significant difference (Figure 2). These results are consistent with our prior work [12].

**Figure 2:**
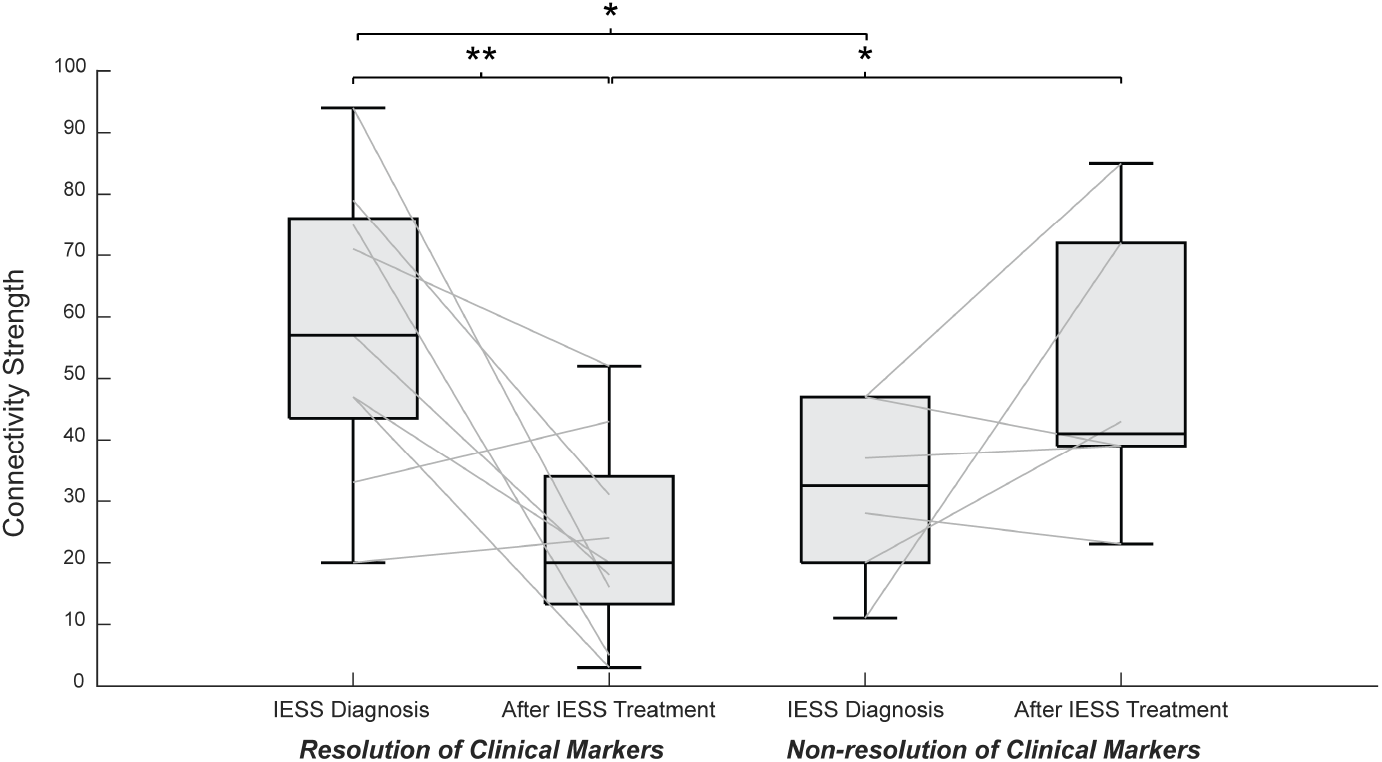
EEG connectivity strength is a candidate biomarker for IESS treatment response. Connectivity strength decreases after IESS treatment for 7/9 subjects with resolution of clinical markers (left) and it increases for 4/6 subjects with non-resolution of clinical markers (right). The change in connectivity strength from the time of IESS diagnosis to the subsequent EEG (after IESS treatment) was significant only for subjects with resolution of clinical markers (Wilcoxon rank sum test, *: p < 0.05; **: p < 0.01).

Across longitudinal data, patients who eventually achieved resolution of SSW exhibited connectivity strength that fluctuated, but generally remained low (Figure 1C), while those that demonstrated persistent SSW had generally increasing connection strength over time (Figure 1D).

Individual subject results also show that changes in connectivity strength frequently reflect disease state/severity (Figure 3). After the diagnosis of IESS, subjects 2 and 3 initially experienced a resolution of epileptic spasms, coinciding with a reduction in connectivity strength during the second and third EEG recordings following diagnosis. Subject 9, however, displayed an increase in connectivity strength following IESS treatment, coinciding with the reappearance of hypsarrhythmia and the persistence of epileptic spasms. The three subjects were then diagnosed with LGS at EEG recordings four, four and five, respectively. After LGS treatment, subject 2 demonstrated resolution of SSW (denoted by an open star) and a notable reduction in connectivity strength. In contrast, subjects 3 and 9 did not respond to treatment (denoted by filled stars), showing similar connectivity strength compared to the time of diagnosis.

**Figure 3:**
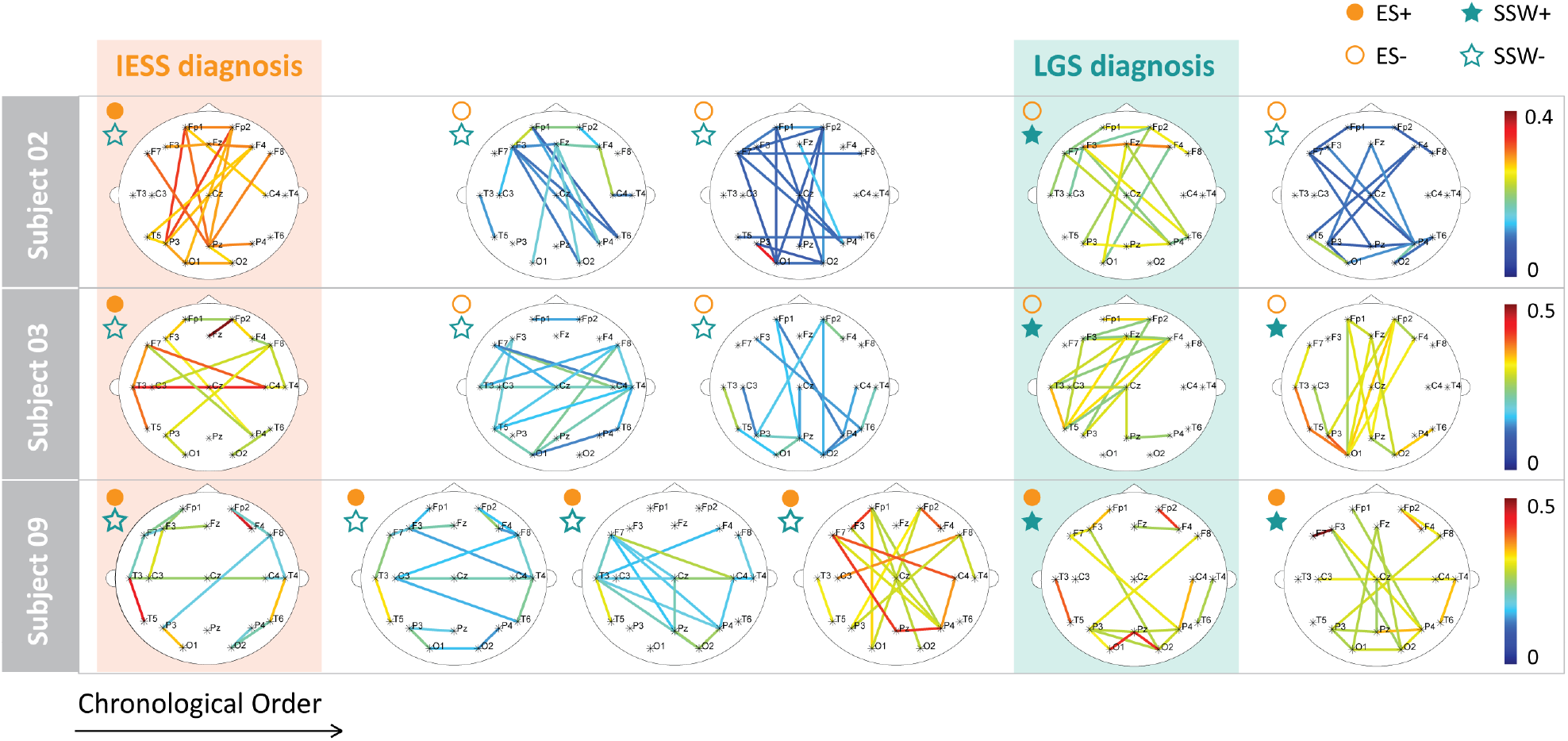
EEG network maps for individual subjects support the potential use of connectivity as a biomarker of diagnosis and treatment response. Here, three individual subjects’ data are shown as examples. Each row corresponds to the topographic connectivity maps of one representative subject, with each network map representing a separate EEG study, chronologically ordered from left to right. Colored rectangles highlight the times of IESS diagnosis (orange) and LGS diagnosis (green); they are aligned for all subjects. The annotation at the top left of each network indicates the presence (filled) or absence (non-filled) of epileptic spasms (circle) or SSW (star). Each network map shows the 10% of connections with the highest strength from the individual study, with the connection strength represented by color, to highlight network structure.

### 3.2. Changes in connectivity strength cannot be explained by any single clinical marker

Connectivity strength was only weakly correlated to IED amplitude and not correlated to age (Figure 4), suggesting that the network differences are not due to normal physiological changes associated with development. We then assessed whether the value of connectivity strength could be explained by standard visual features of the EEG. While connectivity strength was significantly higher for EEGs with hypsarrhythmia, SSW, spasms, and GPFA than EEGs without those markers (Figure 5A-D, Wilcoxon rank sum test, p < 0.001), a more detailed analysis revealed limitations in their ability to predict connectivity strength. A multiple linear regression model was applied to quantify the relationship between the presence or absence of these clinical markers and connectivity strength. SSW (p < 0.001) was the only statistically significant predictor of connectivity strength, and the model explained only 28.4% of the variance in connectivity, with a moderate normalized root mean squared error of 25.1%.

**Figure 4:**
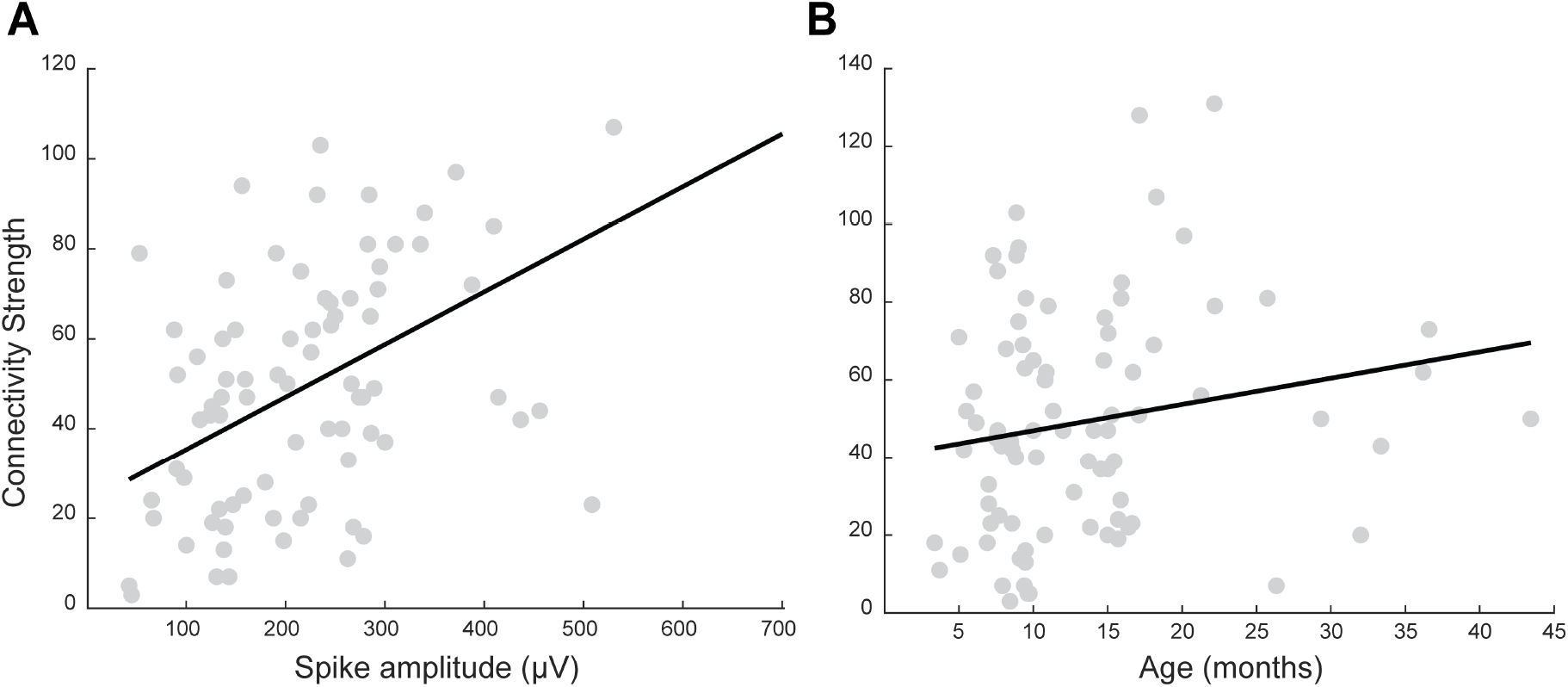
(A) Connectivity strength is weakly correlated to spike amplitude (Pearson correlation, p < 0.001, R2 = 0.30). (B) There is a nonsignificant trend in the correlation between age and connectivity strength (Pearson correlation, p = 0.09, R2 = 0.03).

**Figure 5:**
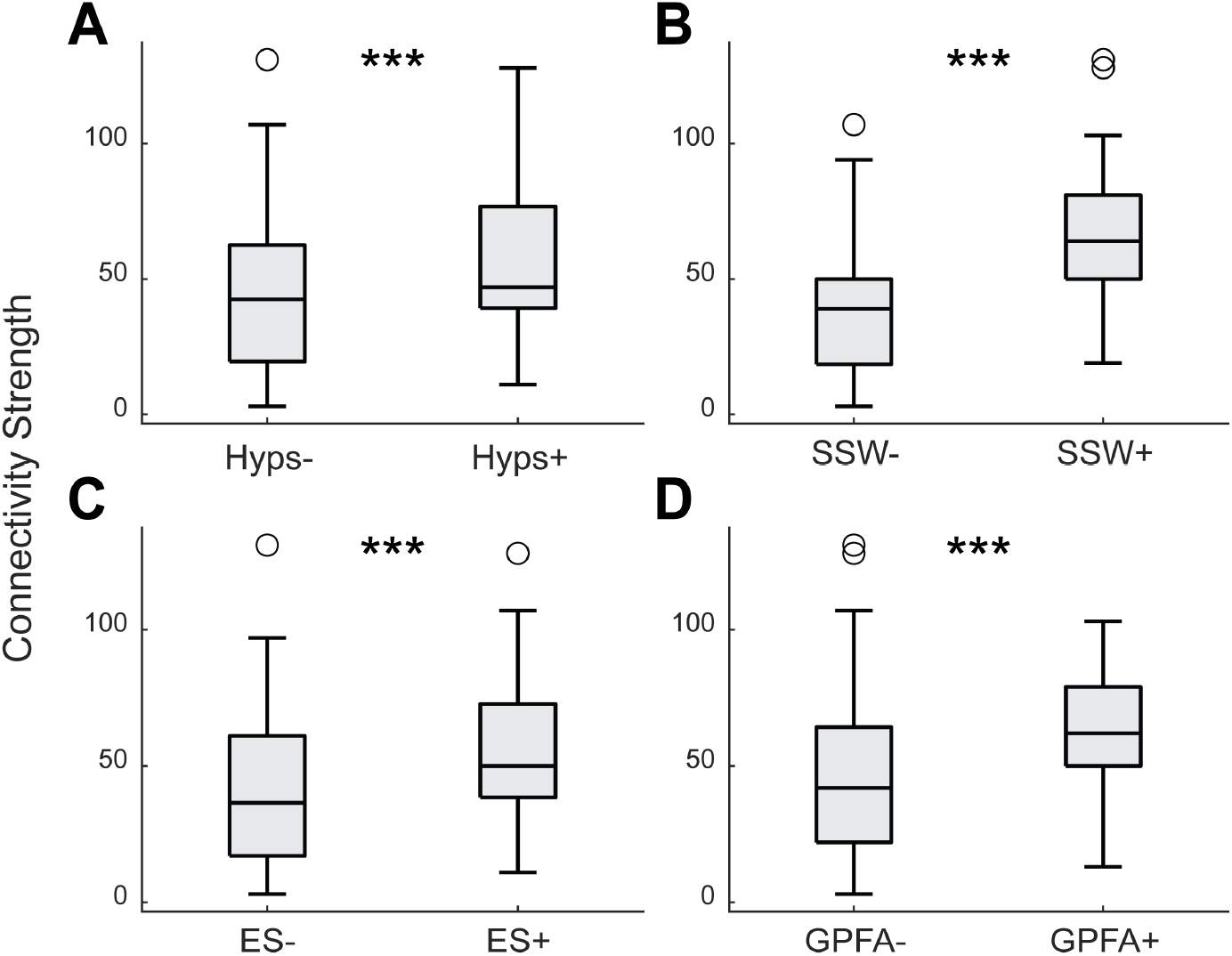
High connectivity strength is associated with several standard clinical EEG markers. Boxplots of connectivity strength are shown for EEGs without (left) and with (right) (A) hypsarrhythmia, (B) SSW, (C) spasms, and (D) GPFA. Wilcoxon rank sum test, ***: p < 0.001.

This suggests that visual features, while associated with increased connectivity, are not strong predictors of connectivity strength. Further, each patient exhibits a different combination of these markers at each point in time; clinically, the impact of each marker cannot be assessed in isolation from the others. For example, in subject 11, the fourth clip shows high connectivity values, with only GPFA present (Figure 1C).

In the fifth clip, there is a slight decrease in connectivity coinciding with the resolution of all clinical markers. Then, the sixth clip (at LGS diagnosis) shows a connectivity level comparable to the fourth clip, with hypsarrhythmia, spasms, SSW and GPFA all present. In the final clip, a notable decrease in connectivity is observed, despite the continued presence of spasms and GPFA.

## 4. Discussion

This study represents a pioneering effort in the development of objective biomarkers to assess the progression from IESS to LGS. To our knowledge, this is the first time that EEG-based FCNs have been proposed as a biomarker during this critical period. Specifically, EEG-based functional connectivity demonstrates promise as a biomarker for the onset of LGS (Figure 1A) and assessment of LGS treatment response (Figure 1C-D). This computational metric appears to have added value beyond the capabilities of traditional clinical markers (Figure 5). Because results are consistent at the single subject level (Figure 3), they could be leveraged to guide clinical management.

Integrating connectivity measures as a standard clinical procedure could significantly enhance both the diagnosis and management of patients. For example, all responders had a connectivity value of 40 or lower in their last EEG (Figure 1C). This threshold could serve as a guideline for assessing treatment response and informing clinical decisions. If connectivity exceeds 40, maintaining or intensifying treatment could be recommended to optimize patient outcomes. Additionally, multiple subjects (e.g., subjects 6, 7, 10, 11, 13) exhibited a dramatic increase in connectivity strength in an EEG clip preceding the LGS diagnosis. Such a finding could alert clinicians of a high likelihood of impending LGS, potentially allowing for an earlier intervention with LGS-specific medications, improving prognosis.

Connectivity strength was related to the presence of standard clinical EEG markers (Figure 5) and weakly correlated to spike amplitude (Figure 4A). The relationship to spike amplitude is consistent with a prior result from our group [18], which demonstrated that only spike amplitudes above 286 µV induce relevant changes in both the network structure and mean connectivity. In the dataset analyzed here, a low proportion of EEG clips contained spike amplitudes of this magnitude, potentially explaining the weak correlation observed between connectivity strength and spike amplitude. While connectivity strength was higher when hypsarrhythmia, ES, SSW, or GPFA was present in the EEG recording, knowledge of these clinical markers was insufficient to predict the connectivity strength of a given EEG. The distributions in Figure 5 have considerable overlap and high variance, making them significant in group level comparisons, but not valuable for predictions about individual subjects. Overall, this suggests that connectivity strength provides additional information that is independent from known clinical EEG markers and can be calculated in an objective manner. Lastly, the lack of a relationship with age (Figure 4B) is consistent with prior studies showing no correlation between connectivity strength and age in IESS patients and controls less than four years old [10] and in healthy infants less than two years old [17].

An important avenue for future work lies in exploring not just the magnitude of the connectivity, but also the structure of the connectivity network, to better understand the progression from IESS to LGS. Previous studies have shown significant differences in connectivity strength between controls and IESS subjects [10-12]. The connectivity networks of control subjects are typically organized into two bilaterally symmetric clusters: (1) occipital and posterior temporal head regions (electrodes T5, T6, O1, and O2), and (2) frontopolar, frontal, and anterior/mid-temporal head regions (electrodes Fp1, Fp2, Fz, F3, F4, F7, F8, T3, and T4). However, IESS subjects exhibit additional strong cross-hemispheric connections between frontal and parietal regions [12]. This difference in network organization between controls and IESS subjects suggests that both connectivity strength and network structure are valuable indicators of disease evolution and treatment response. Future studies could focus on characterizing these networks during the progression from IESS to LGS, ideally while controlling for each patient’s underlying etiology.

There were several limitations to our study. The study sample size was limited to fifteen for this descriptive observational study, as IESS is a rare disease, occurring in 2-5 of 10,000 infants [15], and here we are limited to a subset of that population. Moreover, we required longitudinal EEG data from all patients, which was typically only available if they were treated for both IESS and LGS at the same medical center, further limiting the population size. This enabled us to obtain longitudinal EEG data from each subject, but it also resulted in a relatively small number of subjects with heterogeneous etiologies (Table 1). The retrospective nature of the data was associated with additional limitations. We were not able to standardize the number of EEG recordings or the time intervals at which EEGs were collected. For each subject, EEG studies were done when clinically indicated, which could cause a bias toward abnormal studies. Further, we were unable to directly assess the resolution of LGS and therefore relied on the presence or resolution of SSW activity as a surrogate marker. In addition, robust longitudinal neurocognitive data were not available for our cohort and thus assessing the predictive power of functional networks as a biomarker of long-term outcome was not possible. Lastly, clinical data were to be extracted from the medical record. To mitigate potential bias of clinical EEG interpretation in the medical record, we conducted a blinded review of each EEG to confirm the presence/absence of seizures and other salient EEG features.

The high variability across subjects is both a strength and a limitation of the study. Despite the differences in underlying etiology, age, medications, and course of disease evolution (i.e., whether or not spasms relapsed and whether or not SSW was resolved by treatment), we found robust changes in EEG connectivity strength relative to the IESS and LGS diagnostic timepoints. This demonstrates the potential for broad application of this computational metric and its potential as an objective biomarker. However, we also encountered many challenges in quantifying and standardizing the assignment of clinical markers across this highly variable population. For instance, subject 4 did not exhibit epileptic spasms at the time of IESS diagnosis, as they developed clinical spasms earlier that were not captured during their IESS diagnostic EEG study just prior to treatment initiation. Subjects 5, 10, 11, and 13 exhibited SSW before they were given their LGS diagnoses because they did not develop a second seizure type until a subsequent EEG study. Further, all subjects except for subjects 2, 3, 6, and 12 exhibited epileptic spasms during and/or after LGS diagnosis, showcasing the continuum upon which these diseases reside. This heterogeneity hinders the generalization of results to a broader population, highlighting the need for personalized approaches in both diagnosis and treatment, while also clearly demonstrating the need for objective biomarkers that provide a more continuous measurement of disease severity. The variability in clinical presentations emphasizes the need for EEG-based computational biomarkers to standardize and support the evaluation and decision-making process.

Ultimately, this work suggests EEG functional connectivity as a framework for understanding and accurately predicting the evolution of IESS to LGS. This could facilitate the prevention of LGS via early, effective intervention, thus leading to improved outcomes and quality of life for children at high risk of developing LGS.

## 6. Acknowledgements

This work was supported by a grant from the Lennox-Gastaut Syndrome Foundation, a grant from the Physician-Scientist Scholars program at the Children’s Hospital of Orange County, and the UC Irvine California-Catalonia Engineering Program, through a Balsells Fellowship to BRM.

## 7. Authors’ Contribution

BRM has first authorship of the manuscript and performed data analysis, figure preparation, wrote the first manuscript draft and helped edit the manuscript. RS, DH and NB contributed data analysis. VL, DS, MS, DP, DA, and CS provided clinical expertise, reviewed clinical data, and assessed clinical markers. DS oversaw EEG and clinical data collection and EEG visual assessment, and he assisted in manuscript writing and editing. BL helped conceptualize and supervise the data analysis and manuscript preparation and contributed to writing and editing the manuscript. All authors contributed to the article and approved the submitted version.

